# Plasma Circular RNA Panel to Diagnose Hepatitis B Virus-Related Hepatocellular Carcinoma

**DOI:** 10.1101/576751

**Authors:** Jian Yu, Meng-chao Wang, Wen-bing Ding, Xing-gang Guo, Jian Xu, Qing-guo Xu, Yuan Yang, Shu-han Sun, Jing-feng Liu, Lun-xiu Qin, Hui Liu, Fu Yang, Wei-ping Zhou

**Affiliations:** The Third Department of Hepatic Surgery, Eastern Hepatobiliary Surgery Hospital, Second Military Medical University, Shanghai 200438, China; Key Laboratory of Signaling Regulation and Targeting Therapy of Liver Cancer (SMMU), Ministry of Education. Shanghai 200438, China; Shanghai Key Laboratory of Hepatobiliary Tumor Biology (EHBH), Shanghai 200438, China; Department of Laboratory Diagnosis, Changhai Hospital, Second Military Medical University, Shanghai 200438, China; Department of Medical Genetics, Second Military Medical University, Shanghai 200433, China; Shanghai Key Laboratory of Cell Engineering (14DZ2272300), People’s Republic of China; Mengchao Hepatobiliary Hospital, Fujian Medical University, Fuzhou, 350025, China; Department of General Surgery, Huashan Hospital & Cancer Metastasis Institute & Institutes of Biomedical Sciences, Fudan University, Shanghai 200040, China

**Keywords:** Circular RNA, CircPanel, Diagnosis, Hepatocellular carcinoma, Plasma

## Abstract

To explore whether plasma circular RNAs (circRNAs) can diagnose hepatitis B virus (HBV)-related hepatocellular carcinoma (HCC), microarray and qPCR were used to identify plasma circRNAs that were increased in HCC patients compared with controls (including healthy controls, chronic hepatitis B, HBV-related liver cirrhosis and HCC patients). A logistic regression model was constructed using a training set (n=313) and then validated using another two independent sets (n=306 and 526, respectively). Area under the receiver operating characteristic curve (AUC) was used to evaluate diagnostic accuracy. We identified a plasma circRNA panel (CircPanel) containing three circRNAs (hsa_circ_0000976, hsa_circ_0007750 and hsa_circ_0139897) that could detect HCC. CircPanel showed a higher accuracy than AFP (alpha-fetoprotein) to distinguish individuals with HCC from controls in all three sets (AUC 0.863 [95% CI 0.819–0.907] vs 0.790 [0.738–0.842], P=0.036 in training set; 0.843 [0.796–0.890] vs 0.747 [0.691–0.804], P=0.011 in validation set 1 and 0.864 [0.830–0.898] vs 0.769 [0.728–0.810], P<0.001 in validation set 2). CircPanel also performed well in detecting Small-HCC (solitary, ≤3cm), AFP-negative HCC and AFP-negative Small-HCC.

**Significance of this study:** **What is already known about this subject?**

1. The diagnostic accuracy of alpha-fetoprotein (AFP) in detecting hepatocellular carcinoma (HCC) is unsatisfactory.
2. Circular RNA (circRNA) expression profiles in HCC and adjacent nontumor liver tissues are significantly different.
3. Plasma circRNAs are enriched, stable and can be biomarkers for various diseases.

**What are the new findings?**

1. The expression of circRNAs in the plasma from HCC patients and chronic hepatitis B is significantly different.
2. Plasma circRNA panel (CircPanel, including hsa_circ_0000976, hsa_circ_0007750 and hsa_circ_0139897) has a higher accuracy than AFP to distinguish individuals with HCC or Small-HCC (solitary, ≤3cm) from controls (healthy controls, chronic hepatitis B and HBV-related liver cirrhosis).
3. CircPanel also performs well in diagnosing AFP-negative HCC and AFP-negative Small-HCC.

**How might it impact on clinical practice in the foreseeable future?**

Plasma CircPanel can be a diagnostic biomarker in detecting HCC and improves the diagnostic accuracy.

## Introduction

Hepatocellular carcinoma (HCC), largely attributable to chronic hepatitis B virus (HBV) infection, is the second most common gastrointestinal solid tumors and remains the second leading cause of cancer-related death in China^1^. The high mortality of HCC is due partly to the fact that early-stage HCC shows no obvious symptoms and the diagnostic accuracy of AFP (alpha-fetoprotein, a serum biomarker for the diagnosis of HCC in clinical use) is unsatisfactory. The sensitivity and specificity of high serum AFP for HCC were reported to range from 39–64% and 76–91%, respectively. ^2–4^ Therefore, a novel biomarker for the detection of HCC, especially early-stage HCC, need to be identified.

Circular RNAs (termed circRNAs) are covalently closed, single-stranded and stable transcripts.^5^ In our previous study, we demonstrated that circRNA expression profiles in HCC and adjacent nontumor liver tissues are significantly different and circular RNA cSMARCA5 inhibits the growth and metastasis of HCC.^6^ Furthermore, it has been reported that plasma circRNAs are enriched, stable and can be biomarkers for non-small cell lung cancer and systemic lupus erythematosus^7^. In this study, using microarray and qRT-PCR (quantitative real-time polymerase chain reaction), we tried to explore whether plasma circRNAs can be biomarkers to diagnose HBV-related HCC (referred to below as HCC).

## Patients and Methods

### Study Design and Participants

The study design is listed in Figure 1. In total, 1195 plasma samples, 40 paired HCC and adjacent noncancerous liver (ANL) tissues were collected from three hospitals in China. The recruited participants were defined as healthy individuals, patients with chronic hepatitis B (CHB), patients with HBV-related liver cirrhosis (referred to below as liver cirrhosis), or patients with HCC by medical doctors, according to eligibility criteria listed in Supplementary Table 1.

**Figure 1.**
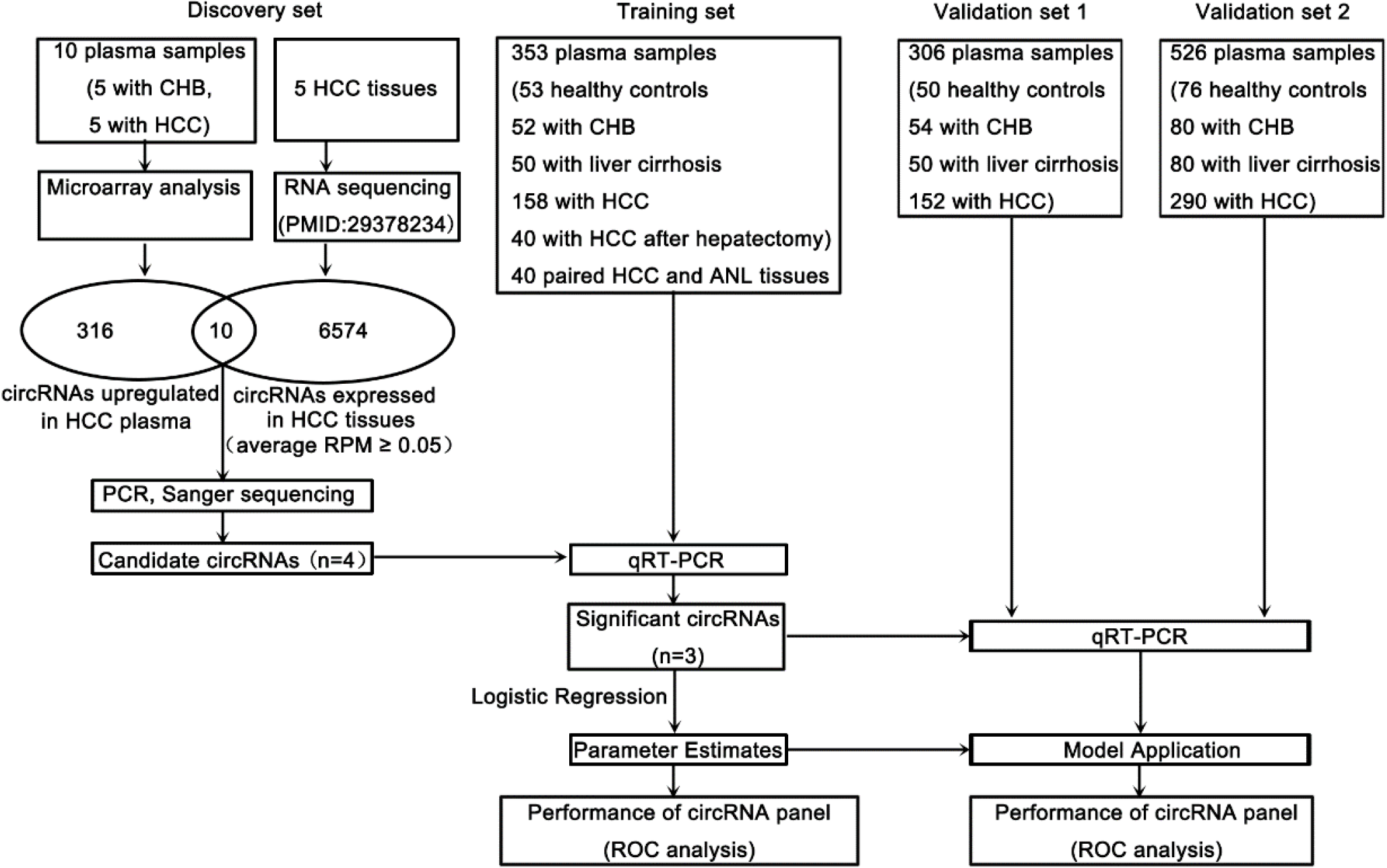
Study design. Abbreviations: ANL, adjacent noncancerous liver; CHB, chronic hepatitis B; HCC, hepatocellular carcinoma; PCR, polymerase chain reaction; qRT-PCR, quantitative reverse-transcriptase polymerase chain reaction; ROC, receiver operating characteristics.

The plasma, HCC and ANL tissues in the discovery and training sets were collected between July 2016 to June 2017 at the Shanghai Eastern Hepatobiliary Surgery Hospital, Second Military Medical University, Shanghai, China. The plasma in validation set 1 was collected between September 2016 to July 2017 at the Shanghai Changhai Hospital, Second Military Medical University, Shanghai, China. The plasma in validation set 2 was collected between February 2016 to May 2018 at the Mengchao Hepatobiliary Surgery Hospital, Fujian Medical University, Fuzhou, China. The details of the clinicopathological characteristics of the participants are listed in Supplementary Table 2. In addition, for the 40 HCC patients undergoing hepatectomy in the training set, we also collected their plasma at the 30th day after hepatectomy, their HCC and paired ANL tissues. The details of the clinicopathological characteristics of these 40 HCC patients are listed in Supplementary Table 3.

Human specimen collection was approved by the ethics committee of each hospital. Written informed consent was obtained from each patient according to the policies of the committee.

### RNA Isolation

For the HCC cell lines and the HCC and ANL tissues, the total RNA was extracted using RNAiso Plus (Takara, Code No. 9109) according to the manufacturer’s instructions.

For the plasma, the total RNA was extracted using the TRIzol™ LS Reagent (ThermoFisher, Code No. 10296010) according to the manufacturer’s instructions. During the isopropanol precipitation, glycogen (catalogue number AM9510, Ambion/Applied Biosystems, Foster City, CA) was added as a coprecipitant (final concentration of 100 μg/mL) to enhance the RNA precipitation.

### CircRNA Microarray Expression Profiling

The total RNAs extracted from the plasma of five HCC patients and five CHB patients were used for microarray analysis as described previously.^8,9^ In brief, the RNAs were digested, amplificated, labelled and hybridized onto the microarray (CapitalBio Technology Human CircRNA Array, Version 2.0). Differential expression analysis of circRNAs was performed using GeneSpring software V13.0 (Agilent). We used threshold values of ≥ 2 or ≤− 2-fold change and a t-test P-value < 0.05. The differentially expressed circRNAs are listed in Supplementary Table 4. The data were Log2 transformed and median centered by genes using the Adjust Data function of CLUSTER 3.0 software and were then further analyzed by hierarchical clustering with average linkage.^10^ Finally, we performed tree visualization by using Java Treeview (Stanford University School of Medicine, Stanford, CA, USA).

### Reverse Transcription

Total RNA from both tissues or plasma was reversely transcribed using M-MLV Reverse Transcriptase Kit (ThermoFisher, Code No. 28025021) according to the manufacturer’s instructions.

### Quantitative Reverse-transcriptase Polymerase Chain Reaction (qRT-PCR)

The qRT-PCR, using SYBR^®^ Premix Ex Taq™ II (Tli RNaseH Plus) and ROX plus (Takara, Code No. RR82LR), was performed on the StepOneTM Real-Time PCR System (Applied Biosystems, Foster City, CA). The PCR primers for β-actin and the four candidate circRNAs (hsa_circ_0000976, hsa_circ_0003506, hsa_circ_0007750 and hsa_circ_0139897) are listed in Supplementary Table 5. The primers for the circRNAs were divergent and circRNA specific. (Supplementary Figure 1A)

For the HCC and ANL tissues, ACTB was employed as the endogenous control, and the relative expression was calculated using the comparative ΔΔCt method.

Since there is no accepted endogenous control for the quantitation of mRNAs/circRNAs in plasma, we used absolute quantitation when detecting the expression of the candidate circRNAs in the plasma as previously described.^11–13^ Briefly, the PCR products were amplified from human pooled plasma cDNA using the primers of the four candidate circRNAs, respectively. Subsequently, the four PCR products were cloned into pUC57 vector, respectively. The resulting constructs were verified by direct sequencing and serially diluted from 5×10^7^ copies/μl to 5 copies/μl, respectively. Those diluted constructs were run in parallel with the samples under identical qPCR conditions and amplified with the same set of primers. A standard curve was generated by plotting the cycle threshold as a function of log10 concentration of the serial diluted controls (Supplementary Figure 1B-E). The relative amount of cDNA of a particular template was extrapolated from the standard curve using the LightCycler software 3.0 (Bio-Rad).

### Statistical Analysis

All statistical analyses were performed using SPSS version 23.0 software (SPSS, Inc., Chicago, IL). For comparisons, chi-squared test, Student’s t test, Wilcoxon signed-rank test, Mann-Whitney U test and Kruskal-Wallis H test were performed, as appropriate. Correlations were measured by Spearman correlation analysis. The optimal cut-off values of the expression of the candidate circRNAs in plasma were determined by a ROC curve (Euclidean distance) analysis in Cutoff Finder^14^ (http://molpath.charite.de/cutoff/). Binary logistic regression was used to build the diagnostic model CircPanel (circRNA panel, including hsa_circ_0000976, hsa_circ_0007750 and hsa_circ_0139897) as described previously.^12,15,16^ Area under the receiver operating characteristic curve (AUC) was used to evaluate diagnostic accuracy. The comparation of AUC was performed by the pROC package of R software (version 3.0.1).^15,17^ All P values were two sided. It was considered to be statistically significant when P<0.05.

**The remaining methods are described in the Supplementary Data 1.**

## Results

### Identification of Circular RNAs By Microarray and PCR in Human Plasma Samples

Using circRNA microarray, we compared the expression of circular RNAs in the plasma from five HCC patients and five CHB patients (discovery set). Among the 371 differentially expressed circRNAs, 326 were upregulated and 45 were downregulated in the plasma from HCC patients compared with that from CHB patients (Supplementary Table 4, Supplementary Figure 2A). We hypothesized that circRNAs in HCC tissues could be secreted into plasma. By overlapping the 326 upregulated plasma circRNAs and the 6584 circRNAs detected in HCC tissues from our group’s previous study^6^, we obtained 10 candidate circRNAs (Supplementary Table 4). Subsequently, we successfully validated four of these (hsa_circ_0000976, hsa_circ_0003506, hsa_circ_0007750 and hsa_circ_0139897) in human plasma (Supplementary Figure 2B-D), human HCC tissues (Supplementary Figure 3A-C), and human HCC cell lines (HepG2 and Huh7) (Supplementary Figure 4) and by qRT-PCR using circRNA-specific divergent primers, agarose gel electrophoresis and Sanger sequencing. Furthermore, we found that incubating plasma at room temperature for up to 24 h had minimal effect on the expression of these four circRNAs (Supplementary Figure 2E), indicating that they were stable in plasma and could be used as a biomarker. In addition, after the treatment with RNase R (a highly processive 3’-to-5’ exoribonuclease that digests linear RNAs^18^), none showed significant changes (Supplementary Figure 3D), which demonstrated that they were truly circular and not linear.

**Figure 2.**
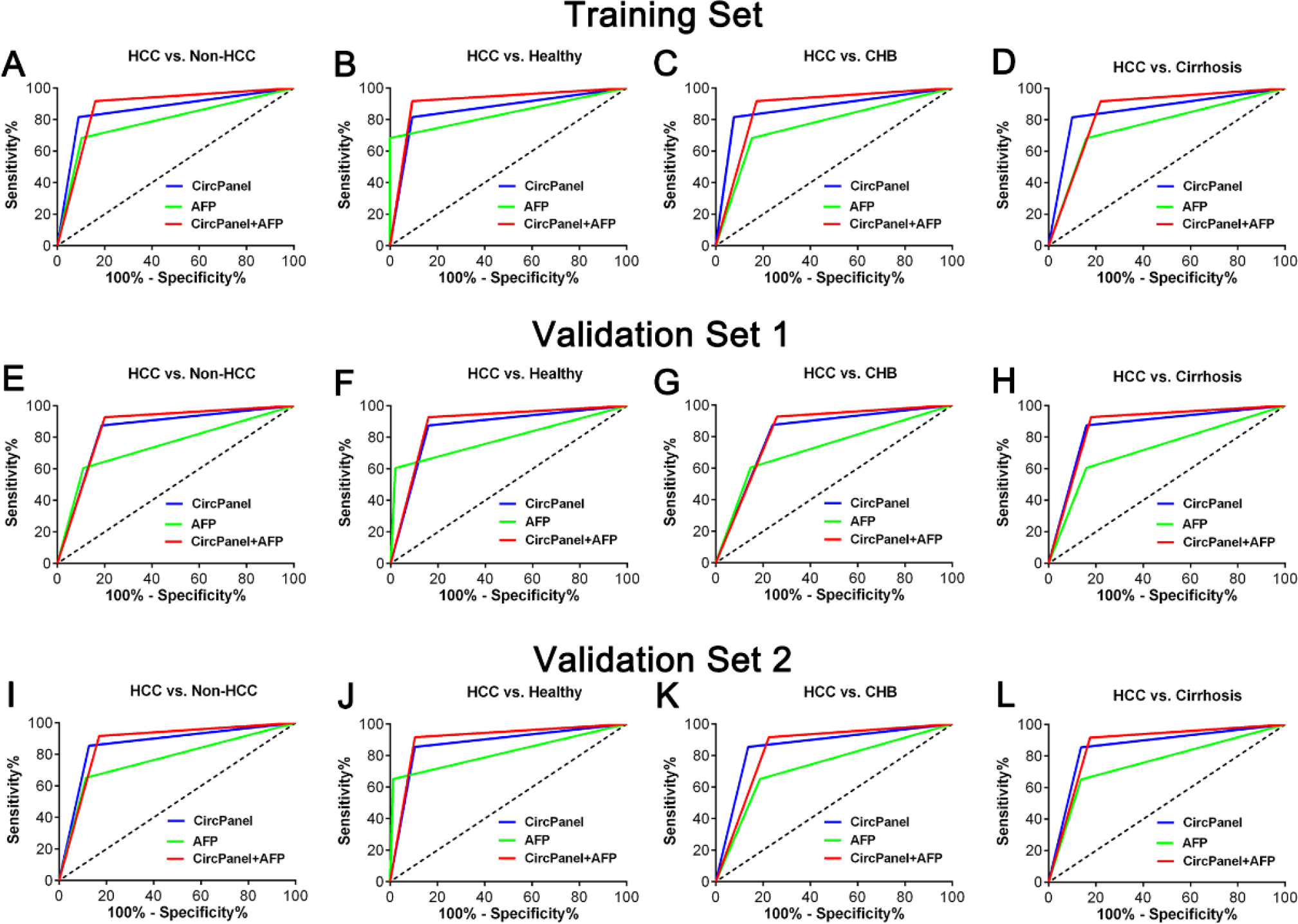
The performance of the CircPanel, AFP and their combination for the diagnosis of HCC in the training set (A-D), validation set 1 (E-H) and validation set 2 (I-L). The detailed diagnostic performances are listed in Table 1. Abbreviations: AFP, alpha fetoprotein; CHB, chronic hepatitis B; CircPanel, circRNA panel containing three circRNAs (hsa_circ_0000976, hsa_circ_0007750 and hsa_circ_0139897); HCC, hepatocellular carcinoma.

**Figure 3.**
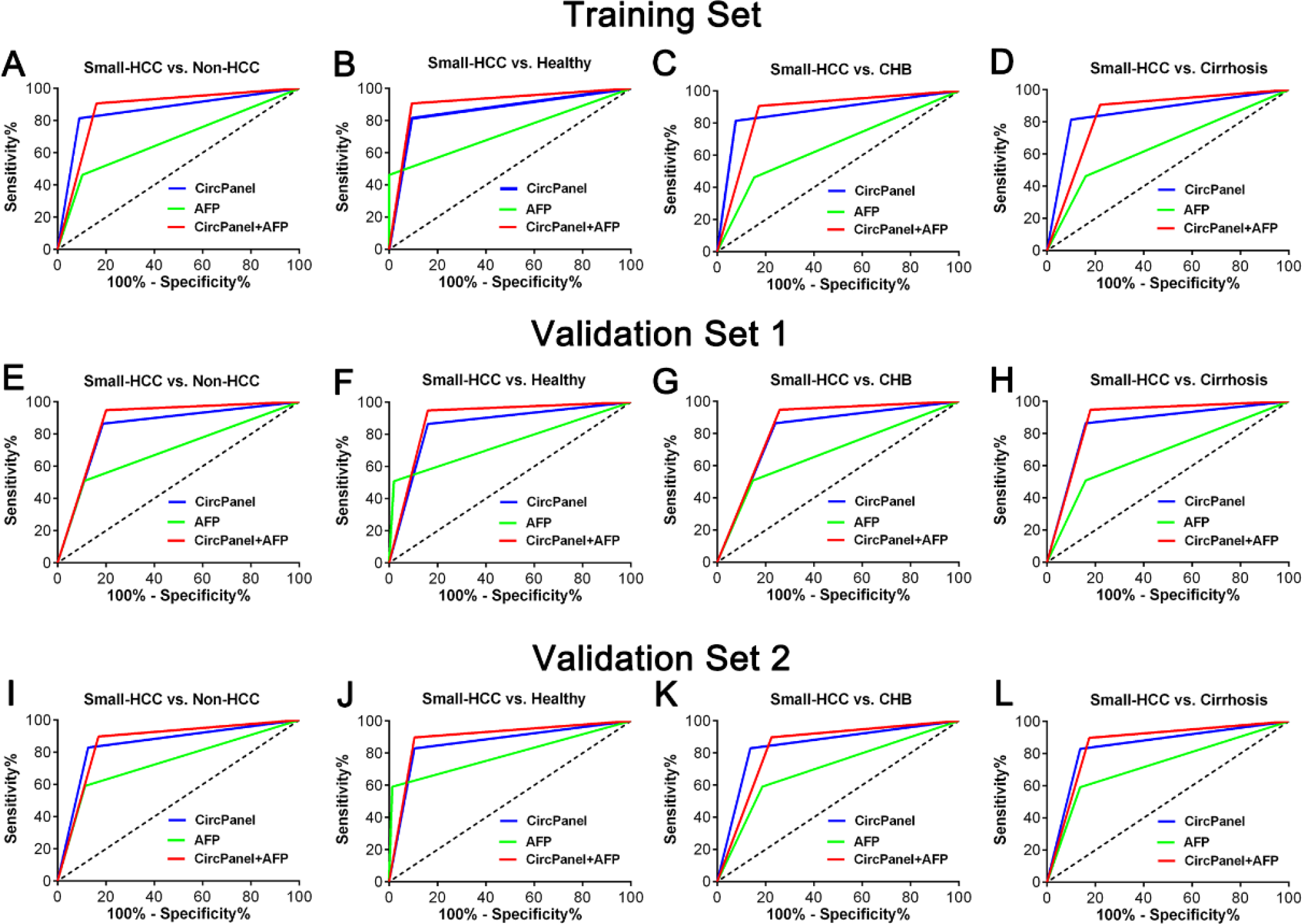
The performance of the CircPanel, AFP and their combination for the diagnosis of Small-HCC in the training set (A-D), validation set 1 (E-H) and validation set 2 (I-L). Thee detailed diagnostic performances are listed in Table 1. Abbreviations: AFP, alpha fetoprotein; CHB, chronic hepatitis B; CircPanel, circRNA panel containing three circRNAs (hsa_circ_0000976, hsa_circ_0007750 and hsa_circ_0139897); HCC, hepatocellular carcinoma; Small-HCC, solitary, diameter ≤3 cm.

Subsequently, HepG2 and Huh7 cells were subcutaneously implanted into the bilateral armpits of BALB/c nude mice. Twenty-eight days later (once tumors were well established), all mice were sacrificed, and their plasma was collected for the detection of the four candidate circRNAs. As expected, agarose gel electrophoresis following RT-PCR (Supplementary Figure 5) and Sanger sequencing (Supplementary Figure 6 showed that these four circRNAs existed in the plasma from the mice in the HepG2 and Huh7 groups but not in the control group. This proved that HCC cells secrete circRNAs into plasma.

### The Building of the Diagnostic Model CircPanel Based on the Training Set

By qRT-PCR, we detected the expression of hsa_circ_0000976, hsa_circ_0003506, hsa_circ_0007750 and hsa_circ_0139897 in the plasma from 158 HCC patients, 53 healthy controls, 52 CHB patients and 50 HBV-induced liver cirrhosis patients (Figure 1) and found that the expression of hsa_circ_0000976, hsa_circ_0007750 and hsa_circ_0139897 (but not hsa_circ_0003506) in the plasma from the HCC patients was higher than that in the plasma from the healthy controls, CHB patients and liver cirrhosis patients (Supplementary Figure 7A). For the 40 HCC patients undergoing hepatectomy in the training set, we also detected the expression of these four circRNAs in their plasma at the 30th day after hepatectomy and in their HCC and paired ANL tissues. We found that the expression of hsa_circ_0000976, hsa_circ_0007750 and hsa_circ_0139897 (but not hsa_circ_0003506) in the plasma from the HCC patients was positively correlated with the expression in their HCC tissues (Supplementary Figure 7B) and was significantly downregulated after hepatectomy (Supplementary Figure 7C). Furthermore, the expression of hsa_circ_0000976 and hsa_circ_0007750 was higher in the HCC tissues than in the ANL tissues (Supplementary Figure 7D). Therefore, we chose hsa_circ_0000976, hsa_circ_0007750 and hsa_circ_0139897 as the candidate circRNAs for the diagnosis of HCC.

Using Cutoff Finder^14^ (http://molpath.charite.de/cutoff/), we determined that the best cutoff values of plasma hsa_circ_0000976, hsa_circ_0007750 and hsa_circ_0139897 for distinguishing HCC and Non-HCC (healthy controls, CHB patients and liver cirrhosis patients as a whole) were 1067, 4324 and 1108 copies/ml of plasma, respectively. Their diagnostic performance is shown in Supplementary Table 6. To improve diagnostic accuracy, using binary logistic regression, we built the diagnostic model CircPanel. The predicted probability of being detected as HCC by the CircPanel was calculated by: Logit (P=HCC)= −3.502+1.920*hsa_circ_0000976+2.800*hsa_circ_0007750+3.154*hsa_circ_0139897. In this equation, the circRNA symbol was substituted with the discretized value one when the level of the circRNA was higher than the corresponding best cutoff value; otherwise, it was substituted with the discretized value of zero. If the result of logit[p=HCC] was higher than 0.5, then the detected sample was predicted as HCC; otherwise it was Non-HCC. As expected, the diagnostic accuracy of the CircPanel was higher than those of hsa_circ_0000976 (AUC 0.863 [0.819–0.907] vs 0.702 [0.644–0.761], P<0.001), hsa_circ_0007750 (AUC 0.863 [0.819–0.907] vs 0.776 [0.723–0.830], P=0.012) and hsa_circ_0139897 (AUC 0.863 [0.819–0.907] vs 0.749 [0.693–0.804], P=0.001) (Supplementary Table 6).

We also analyzed the diagnostic performance of AFP, whose recommended clinical cutoff is 20 ng/ml, in detecting HCC in the training set (Table 1). Furthermore, using the aforementioned method, we combined the CircPanel and AFP (CircPanel+AFP) to diagnosis HCC. The predicted probability of being detected as HCC by CircPanel+AFP was calculated as Logit (P=HCC)=−2.152+3.321*CircPanel+2.241*AFP. The diagnostic performance of CircPanel+AFP was then analyzed (Table 1).

### The Performance of CircPanel, AFP and Their Combination for the Diagnosis of HCC

We then detected the expression of hsa_circ_0000976, hsa_circ_0007750 and hsa_circ_0139897 in the plasma from validation set 1 (152 HCC patients, 50 healthy controls, 54 CHB patients and 50 HBV-induced liver cirrhosis patients) and validation set 2 (290 HCC patients, 76 healthy controls, 80 CHB patients and 80 HBV-induced liver cirrhosis patients) and analyzed the performance of the CircPanel, AFP and their combination (CircPanel+AFP) for the diagnosis of HCC.

As a result, we found that both the CircPanel and CircPanel+AFP showed a higher accuracy than AFP in distinguishing individuals with HCC from Non-HCC in all three sets (CircPanel versus AFP: AUC 0.863 [95% CI 0.819–0.907] vs 0.790 [0.738–0.842], P=0.036 in the training set; 0.843 [0.796–0.890] vs 0.747 [0.691–0.804], P=0.011 in validation set 1 and 0.864 [0.830–0.898] vs 0.769 [0.728–0.810], P<0.001 in validation set 2. CircPanel+AFP versus AFP: 0.878 [0.836–0.920] vs 0.790 [0.738–0.842], P=0.010 in the training set; 0.863 [0.819–0.908] vs 0.747 [0.691–0.804], P=0.002 in validation set 1 and 0.874 [0.840– 0.907] vs 0.769 [0.728–0.810], P<0.001 in validation set 2). In addition, the CircPanel and CircPanel+AFP were not significantly different in distinguishing HCC from Non-HCC (Table 1, Figure 2).

Subsequently, we divided the Non-HCC group into healthy, CHB and liver cirrhosis groups and analyzed the diagnostic performance of CircPanel, AFP and CircPanel+AFP in HCC versus Healthy, HCC versus CHB and HCC versus Cirrhosis. The results were similar, especially for HCC versus CHB and HCC versus Cirrhosis (Table 1, Figure 2).

### The Performance of CircPanel, AFP and Their Combination for the Diagnosis of Small-HCC

We then analyzed the performance of CircPanel, AFP and their combination (CircPanel+AFP) in the diagnosis of Small-HCC (solitary, diameter ≤3 cm) and found that both the CircPanel and CircPanel+AFP showed a higher accuracy than AFP in distinguishing individuals with Small-HCC from Non-HCC in all three sets (CircPanel versus AFP: 0.862 [0.796–0.928] vs 0.680 [0.589–0.770], P=0.001 in the training set; 0.838 [0.776–0.900] vs 0.699 [0.613–0.785], P=0.011 in validation set 1 and 0.851 [0.799–0.903] vs 0.738 [0.671–0.805], P=0.009 in validation set 2. CircPanel+AFP versus AFP: 0.873 [0.817–0.929] vs 0.680 [0.589–0.770], P=0.001 in the training set; 0.874 [0.823–0.925] vs 0.699 [0.613–0.785], P=0.001 in validation set 1 and 0.864 [0.818–0.910] vs 0.738 [0.671–0.805], P=0.002 in validation set 2). In addition, the CircPanel and CircPanel+AFP did not show a significant difference in distinguishing Small-HCC from Non-HCC (Table 2, Figure 3).

Subsequently, we divided the Non-HCC group into healthy, CHB and liver cirrhosis groups and analyzed the diagnostic performance of the CircPanel, AFP and CircPanel+AFP in Small-HCC versus Healthy, Small-HCC versus CHB and Small-HCC versus Cirrhosis. Similar results were obtained, especially in Small-HCC versus CHB and Small-HCC versus Cirrhosis (Table 2, Figure 3).

### The Performance of CircPanel for the Diagnosis of AFP-negative HCC and AFP-negative Small-HCC

Furthermore, we analyzed the performance of the CircPanel in the diagnosis of AFP-negative HCC and AFP-negative Small-HCC. The results showed that the CircPanel also had a high diagnostic accuracy (all AUCs were higher than 0.800, Table 3).

## Discussion

It is reported that plasma circRNA hsa_circ_0001445 is a fairly accurate marker for distinguishing HCC cases from healthy controls as well as liver cirrhosis or CHB patients.^19^ However, this was a single-center study with limited participants (104 HCC patients, 57 cirrhosis patients, 44 CHB patients, and 52 healthy controls).^19^

Our study is unique for the following reasons. First, to our knowledge, this is the first report to compare the expression of circRNAs in the plasma from HCC and CHB patients by microarray. Furthermore, it was a multicenter study with 1155 participants. Importantly, the three circRNAs in the CircPanel proved to be secreted by HCC cells, and their expression in plasma was positively correlated with that in HCC tissues, though the correlation coefficient was relatively low. In addition, the CircPanel showed higher accuracy than AFP in distinguishing individuals with HCC or Small-HCC from the controls and performed well in diagnosing AFP-negative HCC and AFP-negative Small-HCC. All of these findings make the CircPanel a compelling diagnostic biomarker.

There are a few limitations in the present study. First, all of the HCC patients in this study were HBV-related. Further studies are needed to evaluate the performance of the CircPanel in diagnosing HCC caused by other factors. Second, although the expression of hsa_circ_0139897 in HCC and ANL tissues did not show a significant difference, its expression in the plasma from HCC patients was positively correlated with that in their HCC tissues and was significantly downregulated after hepatectomy. The reason for this discrepancy is not clear at the present time and needs further exploration. One possible explanation is that HCC tissues may secrete more hsa_circ_0139897 than ANL tissues. Third, a nested case-control study should be performed to evaluate the diagnostic performance of the plasma CircPanel in detecting preclinical HCC. Furthermore, the follow up of the HCC patients should be continued in order to analyze the relationship between the plasma CircPanel and the prognosis of HCC patients. In addition, since the expression of hsa_circ_0000976 and hsa_circ_0007750 in the HCC tissues was higher than in the ANL tissues, their role in HCC progression awaits further investigation.

In summary, by a microarray screening and qRT-PCR in a multicenter study, we identified a plasma circRNA panel (CircPanel) containing three circRNAs (hsa_circ_0000976, hsa_circ_0007750 and hsa_circ_0139897) that detected HCC. The CircPanel performed better than AFP in diagnosing HCC and Small-HCC and also identified AFP-negative HCC and AFP-negative Small-HCC effectively. Therefore, we believe that the CircPanel can be a potential biomarker in the clinical diagnosis of HCC.

## Authors’ contributions

Conception and design: Wei-ping Zhou, Fu Yang, Hui Liu, Lun-xiu Qin, and Jing-feng Liu

Financial support: Wei-ping Zhou, Fu Yang, Hui Liu and Shu-han Sun

Administrative support: Wei-ping Zhou, Fu Yang and Lun-xiu Qin

Provision of study materials or patients: Yuan Yang, Jian Xu and Jing-feng Liu

Collection and assembly of data: Jian Yu, Meng-chao Wang, Wen-bing Ding, Xing-gang Guo, Jian Xu and Qing-guo Xu

Data analysis and interpretation: Jian Yu, Meng-chao Wang, Wen-bing Ding, Xing-gang Guo, Jian Xu and Lun-xiu Qin

Manuscript writing: All authors

Final approval of manuscript: All authors

## Abbreviations

AFP: alpha-fetoprotein
ALT: alanine aminotransferase
ANL: adjacent noncancerous liver
AUC: area under the receiver operating characteristic curve
BCLC: Barcelona Clinic Liver Cancer
CHB: chronic hepatitis B
CI: confidence interval
CircPanel: circRNA panel containing three circRNAs (hsa_circ_0000976, hsa_circ_0007750 and hsa_circ_0139897)
circRNA: circular RNA
HBsAg: hepatitis B surface antigen
HBV: hepatitis B virus
HCC: hepatocellular carcinoma
PCR: polymerase chain reaction
qRT-PCR: quantitative reverse transcription polymerase chain reaction
ROC: receiver operating characteristics

## Notes

**Conflict of interest:** The authors who have taken part in this study declared that they do not have anything to disclose regarding funding or conflict of interest with respect to this manuscript.

**Financial Support:** This work was supported by the Shanghai Sailing Program (19YF1459600); National Key Research and Development Program of China (2016YFC1302303); National Key Basic Research Program of China (2014CB542102); Science Fund for Creative Research Groups, NSFC, China (81521091); State key infection disease project of China (2018ZX10732202-002-005); National Human Genetic Resources Sharing Service Platform (2005DKA21300); National Natural Science Foundation of China (81372207, 81472689, 81472691, 81502375, 81672345, 81772529); State Key Program of National Natural Science Foundation of China (81330037).

